# Mapping the vRNA Interaction with HIV-1 Integrase

**DOI:** 10.64898/2026.06.01.729371

**Authors:** Jian Sun, Rahul Yadav, Tolga Catmakas, Luke Fisher, Nicholas C. Fitzkee, Jacques J. Kessl

## Abstract

A series of critical interactions within the viral core between the viral RNA (vRNA) and HIV-1 Integrase (IN) has previously been reported. In these studies, contact points between vRNA and IN were identified using RNA-seq and MS-based protein foot-printing approaches. Several IN amino acids located in its C-terminal domain (CTD) were found to be essential for vRNA binding and their alanine substitution severely impacted the correct morphogenesis of the matured viral core. Here, we have extended these studies by performing a comprehensive mapping of the IN-vRNA interaction by deploying RNA crosslinking and NMR methodologies. Together, these approaches were able to identify additional contacts points between the vRNA and IN. Our results reveal several new basic amino acids located in the IN CTD critical for the vRNA-IN interaction, viral replication and correct morphology of the matured viral core.

## 1. Introduction

The Integrase protein (IN) of Human Immunodeficiency Virus type 1 (HIV-1) inserts the reverse transcribed viral DNA (vDNA) into the host chromosome. This catalytic activity of IN, essential for the early stage of the viral replication, has been developed as therapeutic target, and several FDA-approved inhibitors (Raltegravir, Elvitegravir, Dolutegravir, Bictegravir and Cabotegravir)[1] are now commonly used to treat HIV-1 infected individual. The IN is composed of three structurally distinct domains: The N-terminal domain (NTD), the catalytic core domain (CCD) and the C-terminal domain (CTD) [2, 3]. During integration, these three domains work in conjunction to form large and stable multimeric structures called intasomes where vDNA ends are captured by IN [4-8]. The CCD subsequently inserts the vDNA insertion into host chromatin through two sequential magnesium-dependent reactions, 3’-processing and strand transfer. Several mutational studies have revealed that various amino acid substitutions across the IN sequence can influence not only vDNA integration but also other viral replication steps [9, 10]. Consequently, IN mutations have been grouped into two classes: substitutions that selectively impair integration have been assigned to class I mutations and IN mutants that adversely affect multiple steps of virus replication have been grouped as class II [10]. Interestingly, some class II IN mutations can negatively impact correct virus particle maturation and infectivity during the late stage of replication. A hallmark of this phenotype is the mis-localization of the ribonucleoprotein complexes (RNPs) to an eccentric position trapped between the empty capsid (CA) core and the particle membrane in mature virions, whereas normal virions contain RNPs within the CA core. Characterisation of this viral phenotype have been performed either through electron microscopy [11, 12] or sucrose gradient fractionation [12, 13]. Intriguingly, allosteric IN inhibitors (ALLINIs [13-17], which are also known as LEDGINs [18], NCINIs [19], or INLAIs [20]),selectively bind at the IN CCD dimer interface, induce aberrant IN multimerization and consecutively yield non-infectious particles with eccentrically positioned RNPs similar to those seen with several class II IN mutants [11]. Together, these observations strongly suggest that IN not only directs vDNA integration but also contributes to the maturation step during viral egress. Using an array of *in vitro* and *ex vivo* approaches; we and others have previously reported a strong and essential interaction between the viral RNA (vRNA) and HIV-1 IN within the virion [12, 21]. Additionally, these studies [12] have pointed to a significant contribution of the IN CTD. Here, we have performed a comprehensive mapping of the IN-vRNA interaction using both vRNA crosslinking and NMR methodologies. These approaches are able to identify new functionally essential contacts points between the vRNA and IN. Taken together, our results reveal that several basic amino acids located in the IN CTD are critical for the vRNA-IN interaction and the correct morphology of the matured viral core.

## 2. Materials and Methods

### DNA Constructs and Recombinant Proteins

Alanine substitutions were introduced using PCR-site directed mutagenesis (Agilent) and verified by DNA sequencing. For biochemical studies recombinant proteins 6xHis-tagged full length and CTD domain of WT and mutant INs were expressed in *E. coli* and purified by column chromatography[22, 23]. The ^13^C^15^N-CTD protein was expressed in minimal media M9 supplemented with 1g/L ^15^NH4Cl and 2.5g/L ^13^C_6_-glucose.

### IN-RNA Crosslinking for MS/MS

The 57-mer TAR vRNA substrate with a 4-thiouridine (4sU) modification at base 30 was synthesized and obtained from Horizon Discovery (UK). The IN-TAR complex was allowed to form by incubating 4μM IN and TAR vRNA in a 1:1 ratio for 30 minutes at RT in 100 mM NaCl, 1 mM DTT, 0.8 mM CHAPS, 3% glycerol, 25mM HEPES pH 7.4. Crosslinking was performed in ice by subjecting the samples to 30 minutes 365 nm UV irradiation at 600 mJ/cm^2^ (Analytik Jena UVP Crosslinker). Crosslinking efficiency was monitored by either PAG-SDS or western blot using monoclonal IN antibody (NIH AIDS Reagent Program). Crosslinked samples were then incubated with RNase I (Invitrogen) for 30 minutes at 37C, acetone precipitated and Trypsin digested overnight at 37C to yield peptides with RNA fragments of ∼1-2 bases covalently linked.

### Tandem mass spectrometry (MS/MS) analysis

Trypsin digested samples were analysed using electrospray ionization (ESI) MS/MS and RAW data files were processed with the Open search algorithm[24] to identify the crosslinked amino acid residue. As the traditional search engines for modifications require the adduct to be stable during the tandem mass spectrometry (MS/MS) and fragmentation profile of the cross-linked peptides make up complex data, where the adduct mass of the crosslinked species can change during analysis[25], open search algorithm within the MSFragger search engine[26] was utilized. This method provided a comprehensive analysis where every mass shift observed in the MS/MS portion of the data was analysed. Once the adduct identified, isolated MS/MS data peaks were exported via MZmine 3 software[27] and manually confirmed using the ProteinProspector MS-Product tool (http://prospector.ucsf.edu.).

### AlphaScreen-Based RNA Binding Assays

Direct binding of WT or mutant INs to RNA molecules was performed as previously described [12]. Briefly, 50 nM 6xHis-tagged IN was incubated with increasing concentrations of synthetic biotinylated RNAs (Integrated DNA Technologies) at 4C for 1 hr in the AlphaScreen buffer containing 100 mM NaCl, 1 mM MgCl_2_, 1 mM DTT, 1 mg/ml BSA, 25 mM Tris pH 7.4. The final concentration of the nickel and streptavidin AlphaScreen beads (PerkinElmer) in each well was 20 mg/mL. The AlphaScreen signal was recorded on an EnSpire Plate Reader (PerkinElmer) with an AlphaScreen module using the instrument pre-sets. Data were fitted to a Hill equation using Origin software (OriginLab, Inc.).

### NMR and Chemical Shift Perturbation Mapping of CTD-TAR Complex

All HSQC (heteronuclear single quantum coherence) NMR experiments were performed on a 600 MHz Bruker Avance III NMR system equipped with a CP-QCI cryoprobe in 100 mM NaCl, 8 mM CHAPS, 10 mM DTT, 3% glycerol, 25mM HEPES pH 7.4. The amide resonance in 2D ^1^H-^15^N CTD spectra were assigned according to previously reported NMR data[23] and confirmed with 3D measurements for backbone assignment using the double labelled ^15^N-^13^C CTD. Chemical shift perturbations were collected in the presence of an increasing amount of synthetic and unlabelled TAR RNA. After each addition, changes in chemical shifts of the protein resonances were monitored in TROSY ^1^H-^15^N HSQC spectra. A total of four CTD/RNA ratios (150 μM CTD) were examined: 1:0.1, 1:0.2, 1:0.3 and 1:0.4. Experiments were recorded using standard TROSY-resolved Bruker pulse sequences[28] and spectral data were processed using NMRPipe[29] and analysed with NMRFAM-SPARKY[30].

### Viruses and Cells

HIV-1_NL4-3_ molecular clone was used to generate replication competent HIV-1 virions. The mutant viruses were made by introducing the respective substitutions in the IN coding sequence by site directed mutagenesis (Agilent). HEK293T and HeLa TZM-bl cells were grown in Dulbecco’s modified eagle medium (Invitrogen), supplemented with 10% fetal bovine serum (Invitrogen), penicillin (100 I.U/mL) and streptomycin (100 mg/mL) (GIBCO) at 37C and 5% CO_2_.

### Transfection and Virion Preparations

Transfections in HEK293T cells were performed with X-tremeGENE HP (Roche) and the desired plasmid DNA (1:3 for DNA:reagent ratio). 24 hr post-transfection cells were washed, and fresh media was added (with 1mM of saquinavir or DMSO for the virion processing). Cell culture supernatants containing virions were harvested 48 hr post-transfection, filtered through a 0.45 μm filter, and virions were concentrated by ultracentrifugation with a 25% sucrose cushion at 28,000 rpm for 2 hr at 4C in SW41 rotor (Beckman). Isolated virions were assayed by HIV-1 Gag p24 ELISA (ZeptoMetrix) to determine viral production. Levels of infectivity were monitored in TZM-bl reporter cells by assaying luciferase activity with a Luciferase Assay Kit (Promega) [13].

### HIV-1 Viral Core Analyses Using Sucrose Density Gradient Fractionation

For sucrose density gradient fractionation [12, 13] cell-free virions from transfected HEK293T cells were concentrated by ultracentrifugation over a 25% sucrose cushion, pelleted virions were lysed with 0.5% Triton X-100 and were centrifuged through a 30%–70% linear sucrose density gradient. Fractions were collected starting from the top of the gradient and subjected to immunodetection with commercial HIV-1 p24 monoclonal antibody (SinoBiological) to monitor the distribution of HIV-1 capsid.

## 3. Results

### Binding site crosslinking of vRNA TAR construct into full-length IN

WT recombinant full-length IN and a synthetic RNA construct containing a single photoreactive nucleobase were used to pinpoint the orientation and position of the vRNA during viral maturation. While several regions of the viral genome have been shown by CLIP-seq to interact with IN *ex vivo* [12], the vRNA_(1-57)_-trans-activation response element (TAR) of the viral genome was selected as interacting ligand because of: i) its stable and simple structure and ii) it displays high *in vitro* affinity toward IN [12]. For those reasons, TAR has been used in previous studies as *in vitro* surrogate of the IN-vRNA binding activities[31-33]. In this experiment, we have designed and synthetized a TAR substrate that substitute a 4-thiouridine (4sU) at the position 30 (Fig. 1A-star labelled base). This position on the vRNA has been reproducibly identified as primary crosslinking site within the TAR region during CLIP-seq experiments through the T-to-C base changes induced by the reverse transcription step[31]. Previously published K264A/K266A and R269A/K273A IN mutant constructs[31], which have shown to be TAR binding defective, were used here as negative controls (Fig. 1B). Upon binding, 4sU-subtituted TAR covalent crosslinking with full length recombinant IN was initiated by controlled UV irradiation[25]. The formation of the IN-TAR crosslinked adduct was monitored by either PAGE-SDS or western blot (Fig. 1B) and showed a mass shift of +17kDa corresponding to the covalent addition of the RNA construct. This adduct was then subjected to RNase A/T1 treatment and trypsin proteolysis to yield peptides with RNA fragments of ∼1-2 bases covalently linked. Analysis and sequencing of these IN peptides using tandem mass spectrometry (MS/MS) approaches (see Methods section) detected the formation of IN R262 residue crosslinked to the 4-thiouridine at position 30 with an added mass of 128.15 Da (Fig. 1C), corresponding to 4-Thiouracil. Therefore, this sequencing data strongly suggests that within the IN-TAR complex, the nucleotide 30 of TAR is positioned in close proximity to the IN R262 residue as the RNA substrate interacts with several surface exposed and positively charged IN residues (Fig 6).

**Figure 1.**
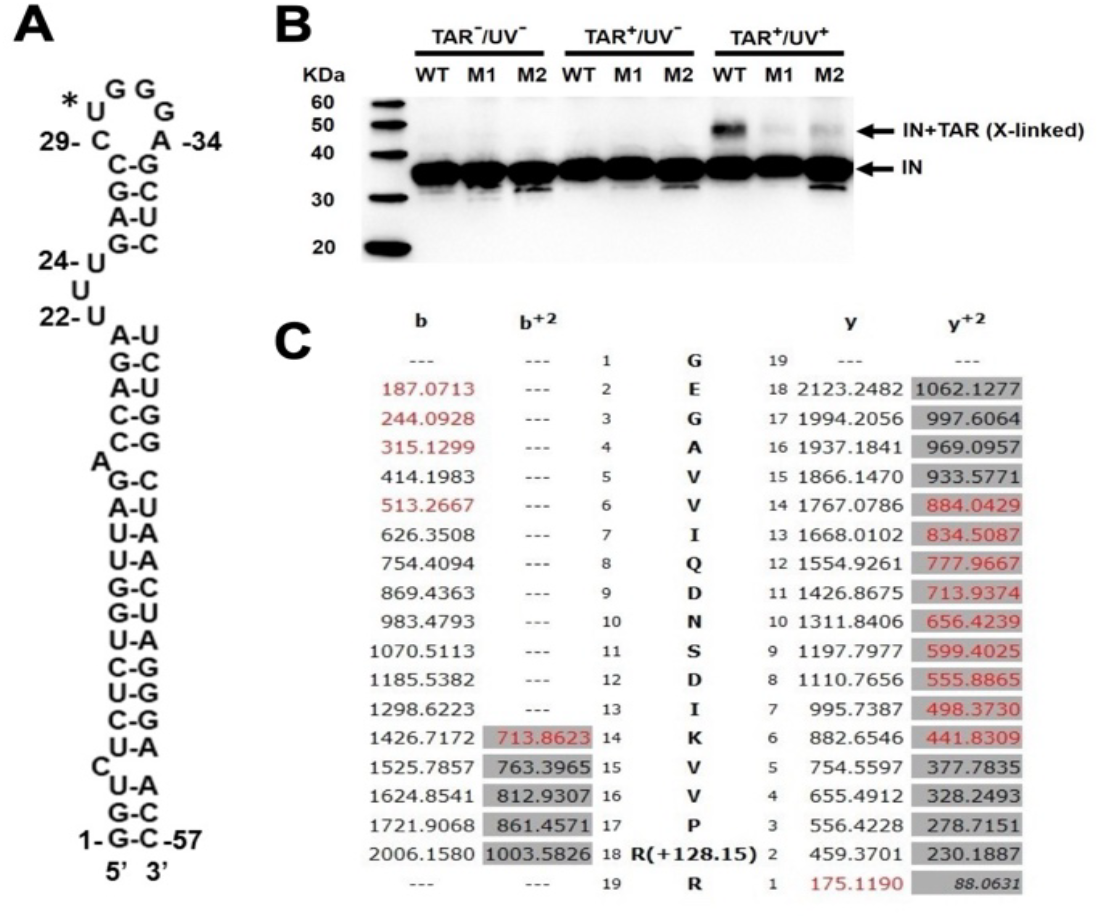
IN-TAR crosslinking. A: TAR sequence with star label on position 30. B: Westerblot of full-length IN proteins after UV-crosslinking experiment using modified 4sU(30)-TAR vRNA construct. The R269A/K273A (M1) and K264A/K266A (M2) IN mutants were used as negative controls. C: MAs spectrometry-based sequencing of the GEGAVVIQDNSDIKVVPRR peptide showing the R262 crosslinked position.

### Mapping TAR binding to IN CTD using protein NMR

As the IN-TAR crosslinking study above and previously published observations[31, 33] suggest, contact points between vRNA and IN point to a major role of the IN CTD. To measure in isolation the specific contribution of this particular IN domain to this interaction, we have used a modified AlphaScreen binding assay, to compare side-by-side the binding affinity of the full length IN and the CTD toward TAR (Fig. 2). While this data confirms a significant binding between CTD and TAR, a lower affinity toward the RNA construct was observed with the IN domain compared to the full-length protein. This 10-folds difference strongly suggests the presence of one or several additional contact points outside the CTD. The binding properties between CTD and TAR were nevertheless compatible with the requirements for NMR detection. As NMR structures of CTD[23] have been previously reported by others, we have used published NMR amide resonance data (Fig. 3A) to assign the obtained signals. In order to map the CTD residues interacting with TAR, we have monitored the chemical shift perturbations (CSP) upon the formation of the CTD-TAR complex. For this we have used a purified ^15^N labeled CTD protein by expressing this construct in minimal media supplemented with ^15^N-ammonium chloride as sole nitrogen source. To measure these CSPs, synthetic unlabeled TAR RNA construct was titrated into the labeled CTD protein while simultaneously acquiring 2D ^1^H/^15^N HSQC (Heteronuclear Single Quantum Coherence) spectrums (Fig. 3B). As several backbone amide resonances shifted upon RNA additions, we were able to pinpoint several surface-exposed CTD residues (Fig. 3B and Table 1) potentially contributing to the interaction with the TAR fragment. NMR resonance signals during the formation of a protein-ligand complex are known to be affected by the rate of exchange between free and bound forms relative to the frequency difference between these states[34, 35]. When the rate is higher (‘fast exchange’), a progressive change in peak position is observed across the titration, while when the rate is lower (‘slow exchange’), separate free and bound resonances are observed with population-dependent intensities. Thus, analyses of exchange rate signals during titration (also referred to as dynamic NMR)[36] were used to reveal binding kinetics on the CTD-TAR complex formation. As high affinity complexes are usually showing “slow exchange” and low affinity complexes in “fast exchange” on the NMR chemical shift timescale[37], different CTD interacting residues were ranked accordingly using rate signal patterns observed during titration (Table 1). The residues Q221, R228 and R263 displayed slow exchange kinetics while F223, K244, R262, K264 and K266 showed fast rates (Table 1).

**Figure 2.**
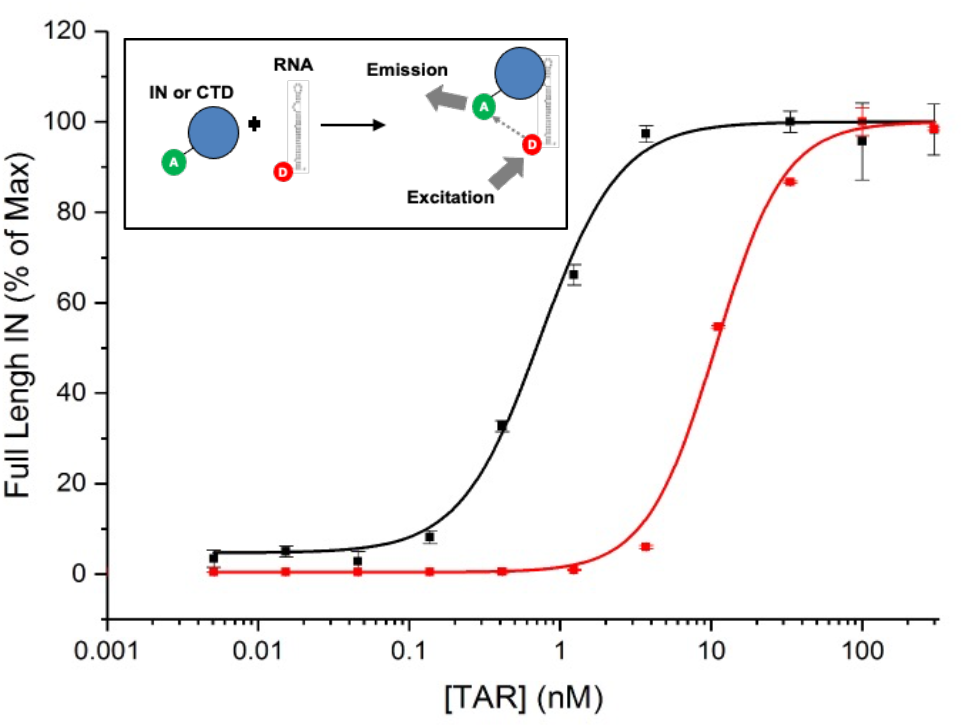
AlphaScreen-based binding affinity measurements of vRNA TAR fragment against full length IN (black) or IN-CTD (red). Assay schematic illustrated in top-left window.

**Figure 3.**
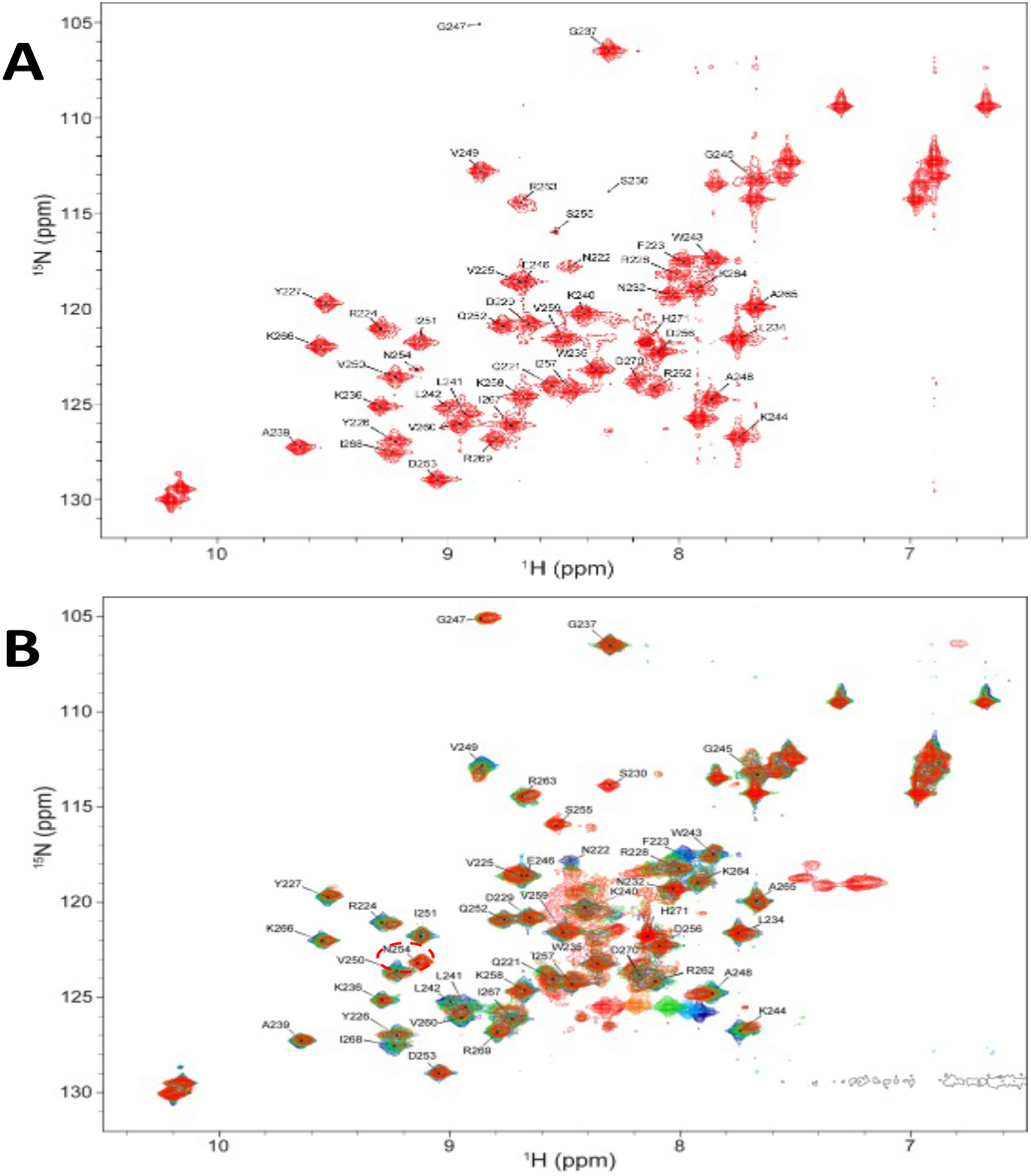
2D Protein NMR of the CTD-TAR complex. A: Signal assignments confirmed by ^15^N/^13^C double labeling experiment with CTD alone. B: Spectrum of ^15^N labeled CTD IN with chemical shift perturbations induced by TAR titration with four CTD/RNA ratios: 1:0.1 (cyan), 1:0.2 (green), 1:0.3 (orange) and 1:0.4 (red).

**Table 1.**
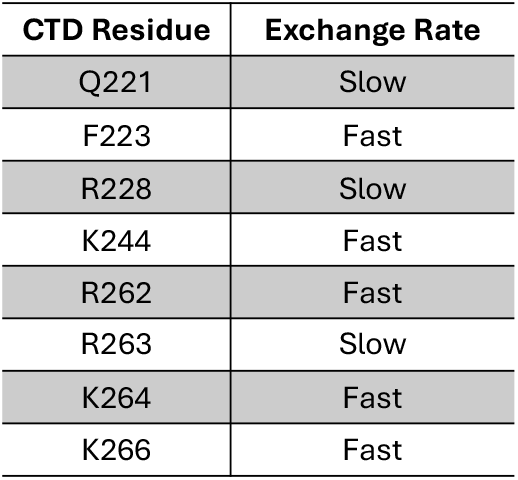
NMR Exchange Rates of identified CTD residues.

| CTD Residue | Exchange Rate |
| --- | --- |
| Q221 | Slow |
| F223 | Fast |
| R228 | Slow |
| K244 | Fast |
| R262 | Fast |
| R263 | Slow |
| K264 | Fast |
| K266 | Fast |

### *In vitro* validation of HIV-1 IN residues interacting with TAR

*In vitro CTD-TAR and IN-TAR binding*. To quantify and validate the contribution of the residues identified above (from both the 4sU crosslinking and the protein NMR experiments) to TAR binding, we have created recombinant mutated CTD and full-length IN proteins with single Alanine substitutions at these specific locations. Each mutated construct was evaluated for their binding affinity toward TAR using the AlphaScreen *in vitro* assay described above (Fig. 2) and compared to the WT proteins. For this, WT or mutated IN proteins (Full length or CTD constructs) were incubated with biotinylated TAR before the addition of AlphaScreen beads. As the K264A/K266A double mutant IN protein has been previously shown to be defective in TAR binding[31], we have used this protein as control. To show the contribution of these 2 residues, we have also introduced the K264A/K266A double mutation into the CTD construct. These *in vitro* measurements revealed that all tested mutations impacted TAR binding (Fig. 4A and 4B). Notably, we observed that the impact of Alanine substitution at each tested position was more severe in the CTD construct (Fig. 4A) than in the context of the full length IN proteins (Fig. 4B) strongly suggesting the contribution of additional contact points with the vRNA construct outside the CTD as measured earlier (Fig. 2).

**Figure 4.**
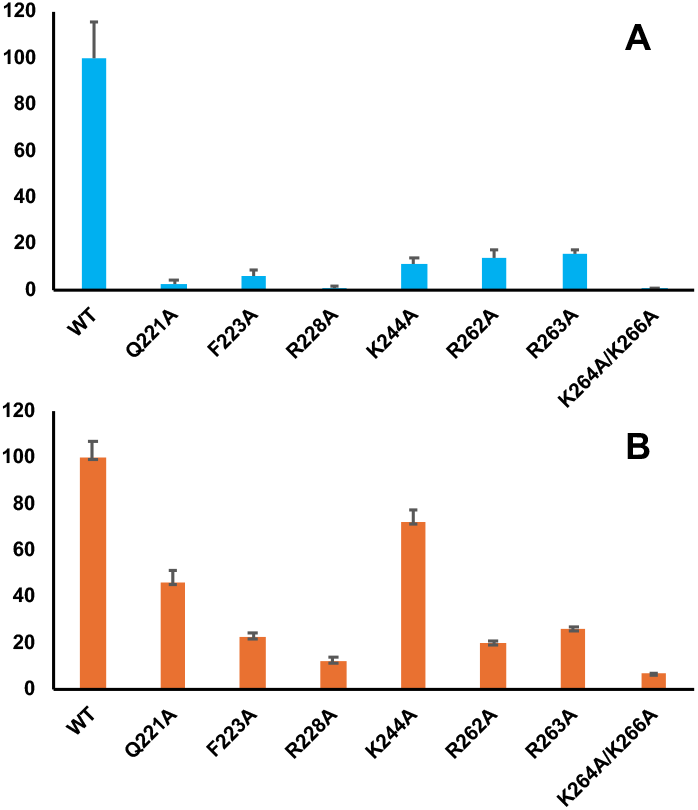
Effect of Alanine substitution on vRNA TAR in vitro binding to full length IN or IN-CTD using AlphaScreen-based assay. A: CTD. B: Full length protein. Values were normalized against WT proteins (100%).

### IN mutations impaired for vRNA binding yield non-infectious virions with eccentric core morphology

To further validate the biological significance of the identified CTD residues, we performed Alanine substitutions at the positions described above in the replication-competent HIV-1_NL4-3_ construct to examine their impact on capsid core morphology and viral replication. Previous studies, including ours and others, have shown that IN mutations affecting vRNA binding influence proper virus particle maturation, leading to the production of non-infectious particles with eccentrically positioned RNPs. This mis-localization of the RNP to a position between the low-density empty capsid core and the particle membrane in mature virions has previously been observed using electron microscopy[9, 21, 33, 38]. Additionally, the buoyant sucrose gradient technique, which measures the relative density of the viral core in mature virions, has also been validated as an effective approach to characterize this mis-localization of the RNP[13, 21, 31, 39]. To confirm the effect of these IN mutations on virus particle maturation, we subjected cell-free and detergent-lysed progeny virions to a linear 20 to 70% (wt/vol) sucrose gradient fractionation. The CA core density was then detected by measuring the HIV-1 CA (p24) content of each sucrose fraction. We used the previously described class II V165A mutation as control as it has consistently been shown to display the eccentric core phenotype[38]. This experiment revealed that specific CTD mutations resulted in a reduction of CA signal in high-density fractions and a simultaneously increase in low density fractions compared to the WT construct (Fig. 5). This observation is consistent with the formation of empty cores due to the mis-localization of the RNPs, characteristic of IN mutants class II phenotype[9]. Among the mutations tested, the F223A exhibited a bimodal distribution of p24 along the gradient (Figure 5) indicating the presence of both eccentric and correct core densities, suggesting a weak or indirect effect on viral morphogenesis. Interestingly, the construct carrying the Q221A mutation was the only one displaying a WT profile, suggesting a marginal impact on viral maturation. To further observe the impact of these IN mutations on viral replication, the infectivity of the virions isolated from transfected cells was measured in target cells and compared to the WT construct (Fig. 6). Except Q221A that retained around 60% of WT infectivity, all considered mutation had a decrease below 30%. Together, these experiments (Figs. 5 and 6) confirmed that single Alanine substitutions at F223, R228, K244 and R263 positions in a replication-competent HIV-1_NL4-3_ construct significantly impact capsid core morphology and viral replication.

**Figure 5.**
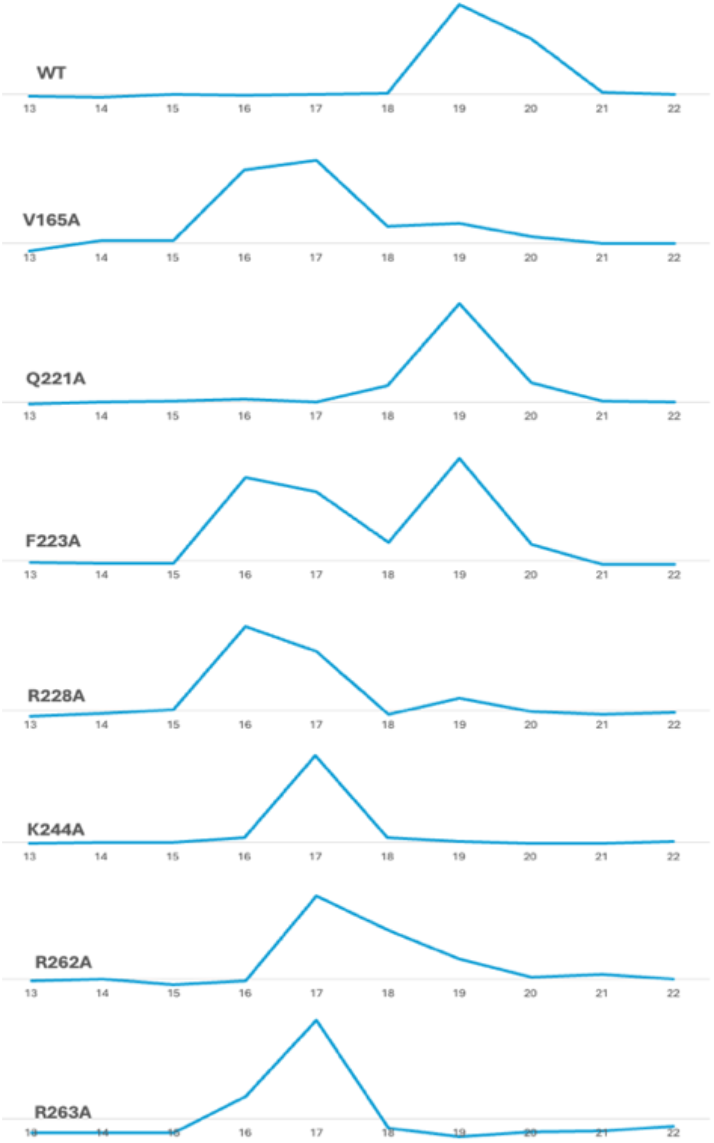
Viral core density profiles of IN WT and mutant virions observed by sucrose fractionation of detergent-lysed virions. The relative capsid (p24) distribution in the bottom fractions is represented.

**Figure 6.**
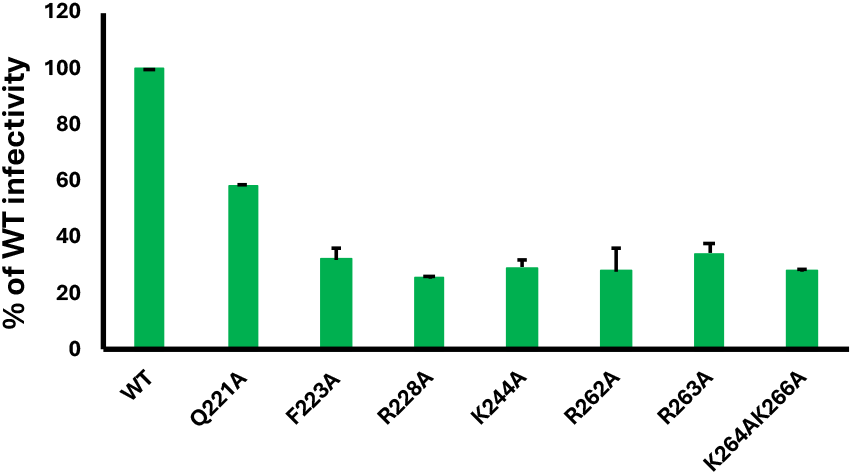
IN mutant virions infectivity measured in target cells. Values were normalized against WT (100%).

## 4. Discussion

Given its involvement observed at multiple stages of viral replication, the HIV-1 IN C-terminal domain (CTD) has been described as the virus’s “Swiss army knife”[40]. With its SH3-like beta-barrel fold structure, this sub-domain appears to perform a wide variety of functions. In early studies, the domain characterized by the presence of several basic residues, has shown substantial non-specific DNA-binding properties [41-43]. Recent intasome structures have pointed to the side chains of IN residues R228, K244, R262, R263, K264, K266 and R269 as components of a basic surface patch capable if interacting with vDNA[40]. A docking study supported by biochemical analysis suggests that INI1, a subunit of the cellular SWI/SNF chromatin remodelling complex and incorporated into viral particles, also interacts with CTD[33]. Early studies have shown that various mutations within IN (class-II IN mutants)[9, 10, 44] could trigger the formation of a particular viral morphological defect in which the RNP condensates outside the capsid core (also described as the eccentric core phenotype). Notably, several of these IN mutants had their mutations located in the CTD[44].

These early observations strongly suggested that IN not only directs vDNA integration but also play a functional role during viral maturation. More recently, IN has been observed as interacting directly with the vRNA during viral egress[31]. The interaction occurs at specific and reproducible sites on the viral genome, mostly on structured elements such as TAR. Preventing this interaction with vRNA by mutating IN, causes the morphological defects described above.

In the present study, we have used a two-pronged *in vitro*-based strategy to probe and map the IN-vRNA molecular interface. We first used full length recombinant IN and a high affinity synthetic TAR 57-nucleotides construct that displays a 4sU modification at position 30[31] to specifically crosslink and identify the IN residue located in the immediate vicinity of this nucleotide. After RNase treatment and trypsin proteolysis of the full-length protein, mass spectrometry-based sequencing reproducibly identified IN R262 residue as likely positioned adjacent to the nucleotide 30 of TAR (Fig. 6, magenta position). Notably, no other crosslinked IN residues were detected. As this IN-TAR crosslinking study above and previously published observations[31, 33, 44] suggest, contact points between vRNA and IN point to a major contribution of the IN CTD.

Because the NMR structure of CTD[23] have been previously reported, we have used this approach to identify additional RNA interacting CTD residues by monitoring chemical shift perturbations (CSP) upon the formation of the CTD-TAR complex. Using this method, we were able to pinpoint eight surface-exposed CTD residues (Q221, F223, R228, K244, R262, R263, K264 and K266) potentially contributing to the interaction with the TAR fragment. Several of those IN residues (R228, K244 and K266) have been previously observed exhibiting class II phenotypes when mutated to Alanine[44]. Interestingly, as discussed above, some of those IN residues (R228, K244, R262, R263, K264 and K266) have also been described as components of a basic surface patch contributing to the CTD interaction with vDNA during integration[40]. The contribution to the TAR binding by each position identified in this study (Q221, F223, R228, K244, R262, R263, K264, and K266) was measured *in vitro* using recombinant proteins (Fig 4) and *ex vivo* in HIV-1_NL4-3_ constructs to examine their impact on capsid core morphology and viral replication. While Alanine replacement in the CTD construct severely impacted TAR binding at all positions (Fig. 4A), the effect was mitigated for Q221A and K244A when full-length proteins were tested (Fig. 4B). Additional *ex vivo* studies confirmed that the infectivity of virions exhibiting the Q221A mutation was marginally impacted with no observed defect on viral core morphology. On the other hand, virions with the K244A IN mutation showed poor infectivity and capsid core density defect, confirming a possible direct contribution to vRNA binding. Interestingly, while other mutated virions (R228A, K244A, R262A and R263A) were exhibiting complete shift toward low viral core densities, the F223A mutant was mixed suggesting a weaker contribution.

While our study was in late writing stage, a new approach exploring the possible morphology of the IN-RNA complex was published[45]. In this work, cryogenic electron microcopy (cryo-EM) was used to model the interactions between IN, vRNA and the capsid core. Because of the unsurmountable challenges cause by the spontaneous aggregation properties of HIV-1 IN at the concentrations needed with this approach, the authors substituted the HIV-1 protein by the cryo-EM compatible SIVtal IN (55% identity with HIV-1 IN). Observation by cryo-EM of concentrated SIVtal IN protein vitrified in the presence of HIV RNA^TAR^ revealed the formation elongated polymers assembled from protein octamers formed around multiple extended duplexes of RNA molecules instead of distinct hairpin-like TAR structures. While the authors acknowledged that the RNA duplex arrangements may have been caused by the specific experimental conditions required for cryo-EM and unlikely to occur during maturation with the full viral genome, several CTD contact points observed with this approach (R228, K244, R262, R263, K264) confirmed our above finding.

Using the data collected in this work, we have generated an energy-minimized molecular model of the CTD-TAR complex (Fig. 7) that locates the crosslinked nucleotide U30 near R262 (Fig. 7, magenta residue) and predicts the relative positions of the RNA fragment according to the NMR shifts (Fig. 7, yellow positions). Interestingly, the proposed model shows the TAR hairpin loop wrapping tightly around a positively charged post structure composed of R262 and R263. Alanine replacement at those positions either individually (this work) or in combination[21, 44] severely impacts RNA binding, infectivity and virion morphology.

**Figure 7.**
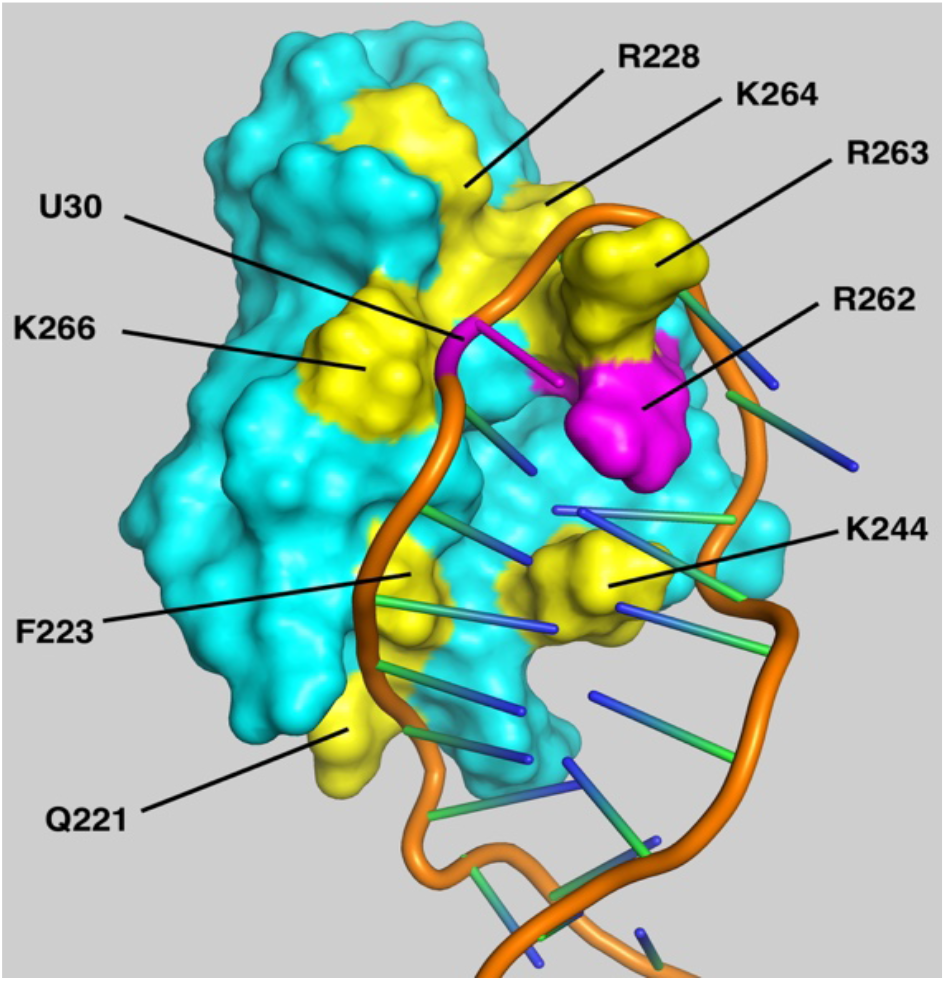
3D Binding Model of the CTD-TAR Complex.

On note, both Q221 and K244 are predicted (Fig. 7) to interact with the ascending and descending stem of the TAR element, which could explain why NMR and the *in vitro* measurements with the truncated CTD protein may artificially amplify the binding contribution of these two residues compared to the full-length protein. More contact points are expected to occur outside the CTD construct used in this work. Most notably is the role of K34 within the IN NTD which has been shown to contribute to vRNA binding[21]. By providing a new experimental-based model of the CTD-TAR complex (Fig. 7), our study offers a molecular basis to previously observed class II mutation phenotypes such as R228A[21], double mutants R262A/R263A[21] and K264A/K266A[31].

## Author Contributions

J.S., R.Y., T.C., L.F., N.C.F. and J.J.K. designed and executed experiments and interpreted results. J.S., R.Y., T.C., and J.J.K. designed the figures. J.S., N.C.F. and J.J.K. wrote the manuscript with constructive input from all authors. All authors have read and agreed to the published version of the manuscript.”

## Abbreviations

The following abbreviations are used in this manuscript:

HIV-1: Human Immunodeficiency Virus type 1
IN: Integrase
vRNA: Viral RNA
TAR: Trans-activation response element
CTD: C-terminal domain
4sU: 4-Thiouridine
CA: Capsid
RNP: Ribonucleoprotein

